# A diffusive homeostatic signal maintains neural heterogeneity and responsiveness in cortical networks

**DOI:** 10.1101/011957

**Authors:** Yann Sweeney, Jeanette Hellgren Kotaleski, Matthias H. Hennig

## Abstract

Gaseous neurotransmitters such as nitric oxide (NO) provide a unique and often overlooked mechanism for neurons to communicate through diffusion within a network, independent of synaptic connectivity. NO provides homeostatic control of intrinsic excitability. Here we conduct a theoretical investigation of the distinguishing roles of NO-mediated diffusive homeostasis in comparison with canonical non-diffusive homeostasis in cortical networks. We find that both forms of homeostasis provide a robust mechanism for maintaining stable activity following perturbations. However, the resulting networks differ, with diffusive homeostasis maintaining substantial heterogeneity in activity levels of individual neurons, a feature disrupted in networks with non-diffusive homeostasis. This results in networks capable of representing input heterogeneity, and linearly responding over a broader range of inputs than those undergoing non-diffusive homeostasis. We further show that these properties are preserved when homeostatic and Hebbian plasticity are combined. These results suggest a mechanism for dynamically maintaining neural heterogeneity, and expose computational advantages of non-local homeostatic processes.

**Author Summary:** Neural firing rates must be maintained within a stable range in the face of ongoing fluctuations in synaptic connectivity. Existing cortical network models achieve this through various homeostatic mechanisms which constrain the excitability of individual neurons according to their recent activity. Here, we propose a new mechanism, *diffusive homeostasis*, in which neural excitability is modulated by nitric oxide, a gas which can flow freely across cell membranes. Information about a neurons’ firing rate can be carried by nitric oxide, meaning that an individual neurons’ excitability is affected by neighboring neurons’ firing rates as well as its own. We find that this allows a neuron to deviate from the target population activity, as its neighbors will counteract this deviation, thus maintaining stable average activity. This form of neural heterogeneity is more flexible than assigning different target firing rates to individual neurons.

## Introduction

Nitric oxide (NO) is a diffusive neurotransmitter which is widely synthesized in the central nervous system, from the retina to the hippocampus [1, 2]. Its properties as a small nonpolar gas molecule allows rapid and unconstrained diffusion across cell membranes, a phenomenon often called *volume transmission* [3]. An important role of NO signaling is to regulate neural excitability through the modulation of potassium conductances in an activity-dependent manner, effectively mediating a form of homeostatic intrinsic plasticity (HIP). Experiments characterizing this effect also demonstrated that NO-synthesizing neurons can induce changes in the excitability of neurons located up to 100 *µ*m away [4, 5]. These findings are corroborated by a recent study demonstrating neurovascular coupling mediated through activity-dependent NO diffusion [6]. We build upon these observations, postulating a general form of HIP mediated by a diffusive neurotransmitter such as NO which we will refer to as *diffusive homeostasis*. This contrasts with canonical models of HIP, here referred to as *non-diffusive homeostasis*, which assume that each neuron has access to only its own activity [7].

Theoretical studies of HIP have generally focused on its role in maintaining stable network dynamics [8, 9]. It has also been recently demonstrated that HIP can improve the computational performance of recurrent networks by increasing the complexity of network dynamics [10]. However, little is known about the effects of HIP on the heterogeneity typically observed in cortical networks; in particular, a growing body of evidence supports the finding that even neurons of the same type have a broad and heavy-tailed distribution of firing rates [11]. Rather than an epiphenomenon of biological noise, neural heterogeneity has been proposed to improve stimulus encoding by broadening the range of population responses [12, 13] However, this form of heterogeneity is difficult to reconcile with canonical models of HIP, which generally suppress cell-to-cell variability [14]. While some degree of heterogeneity in populations of the same type of neuron may emerge naturally [15], we found that such independent sources of variability will generally limit the responsiveness of a network through neuronal saturation.

Using network models and dynamic mean field analysis, here we show that networks with HIP mediated by diffusive neurotransmission exhibit a very different and unexpected behavior. Firstly, we report that diffusive homeostasis provides a natural substrate for flexibly maintaining substantial heterogeneity across a network. Secondly, the resulting population heterogeneity enables linear network responses over a wide range of inputs. This not only improves population coding, but enables a good use of available resources by ensuring that all neurons remain functionally responsive to changes in network dynamics. Finally, we demonstrate that these effects are preserved in networks whose recurrent synaptic inputs undergo Hebbian plasticity.

## Results

We investigated the effects of diffusive homeostasis in a recurrent neural network (Fig. 1A, see Methods) with sparse and random connectivity, based on conventional models of cortical networks giving rise to asynchronous irregular spiking activity [16]. Each neuron received external input with a rate randomly drawn from a normal distribution. Ca^2+^ influx during a spike triggered NO synthesis through nNOS activation (Fig. 1B, see Methods). To simulate spatial NO signaling, neurons were randomly distributed on the surface of a torus and linear diffusion was simulated on this surface. Each neuron’s firing threshold *θ*_*i*_ underwent modulation through negative feedback mediated by the concentration of NO (Equation 6 in Methods).

The effect of a diffusive neurotransmitter mediating HIP within the network was investigated by comparing two cases: first where NO was allowed to diffuse freely across cell membranes as observed experimentally [4], and second without diffusion such that intracellular NO concentration was affected only by a neuron’s own recent activity (Fig. 2). The latter corresponds to a canonical model of HIP as investigated before [8, 9].

### Diffusive homeostasis enables a broad firing rate distribution

Fig. 1C illustrates that both forms of homeostasis stabilized network activity following an increase in input. There was however a crucial difference in how the neurons reacted to this change. While for non-diffusive homeostasis each neuron simply returned to its target firing rate, diffusive homeostasis caused each neuron to sense a mixture of its own activity level and that of the rest of the network. This can be seen in the spatial concentration profiles in Fig. 1C. It is important to note that the spatial position of each neuron was random and independent of its connections, meaning that there was no explicitly defined structure in the NO concentrations.

As a result, these networks exhibited a very different steady state behavior. The firing rate distribution was narrow as expected for non-diffusive homeostasis, but broad and heavy-tailed for diffusive homeostasis (Fig. 1D). The latter is consistent with recent experimental results indicating that firing rate distributions in cortex are generally heavy-tailed, approximating log-normal distributions [11]. There were no noticeable differences in inter-spike interval statistics between networks with diffusive and non-diffusive homeostasis (not illustrated).

We investigated the difference in firing rate distributions by modeling the relation between activity read-out and homeostatic compensation in these two cases using a dynamic mean-field model (see Methods). This approach considered an unconnected population of neurons with random inputs, where each of the two scenarios was simulated by using an appropriate activity read-out. HIP was implemented as in the full spiking model, but the degree of diffusive signaling was now controlled by a single parameter, *α* (Equation 11 in Methods), which determined the balance between local and global activity read-out. If small, neurons used primarily their own activity to modulate their firing threshold, while increasing *α* caused the firing threshold to depend more strongly on the average population activity. Setting, for instance, *α* = 0.8 led to a broad and heavy-tailed rate distribution similar to the full model, while *α* = 0 yielded a narrow distribution as in the non-diffusive case (Fig. 1E).

This model provides a simple and intuitive explanation for this effect. For a non-interacting population, non-diffusive homeostasis can be thought of as precisely matching a neuron’s input *µ*_*i*_ and its threshold *θ*_*i*_ to maintain the target firing rate. We can imitate this by introducing a covariance *σ*(*µ, θ*) between *µ*_*i*_ and *θ*_*i*_, such that a high input rate implies a high firing threshold and a low input rate a low firing threshold. Since setting *α* > 0 (analogous to diffusive homeostasis) introduces a correlation between a neuron’s threshold *θ*_*i*_ and the average population threshold 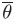, this effectively results in a decorrelation of *µ*_*i*_ and *θ*_*i*_ in comparison with setting *α* = 0 (analogous to non-diffusive homeostasis). In line with the previous results, populations with for instance *σ*(*µ, θ*) = 0.6 yielded a broader and more heavy-tailed distribution of firing rates than populations with *σ*(*µ, θ*) = 0.99 (Fig. 1F).

Since non-diffusive homeostasis directly relates the firing threshold of a neuron to its input, we observed a wider distribution of firing thresholds, which in turn ensured that all neurons assumed similar firing rates. Diffusive homeostasis, on the other hand, yielded similar firing thresholds across the population (Figs. 3A-B). When combined with the nonlinear input-output relation of neurons [17], this gave rise to the broad firing rate distributions we observed (see also Discussion). This result was robust to changes in the rate of NO diffusion. While decreasing the rate of diffusion, *D*, did result in slightly narrower firing rate distributions, they were broader than in networks with non-diffusive homeostasis across a wide range of values (Fig. 3C). A similar trend was observed when varying the width of the external input rate distribution. While decreasing this width led to a decrease in the width of the firing rate distribution, they were consistently broader in networks with diffusive homeostasis (Fig. 3D).

Since one may argue that diffusive homeostasis is merely adding variability to each neuron’s homeostatic signal due to the influence of neighboring neurons’ activity, we now ask whether it is possible to achieve broad firing rate distributions with non-diffusive homeostasis. Indeed, by introducing variability of homeostatic targets (see Methods), we could produce a distribution of firing rates similar to that observed with diffusive homeostasis (Fig. 1D, red histogram). However, as we will show next, the effect of diffusive homeostasis is quite distinct from that of activity-independent, ‘quenched’ heterogeneity arising from randomly distributed homeostatic targets.

### Diffusive homeostasis retains input heterogeneity

To investigate the functional consequences of heterogeneity caused by a diffusive homeostatic process, we next simulated specific changes in external input. First, we stimulated small random groups of neurons at higher input rates of 5Hz and 10Hz (versus a baseline of 2Hz), as illustrated in Fig. 4. Such inputs may, for instance, reflect developmental or other plastic changes that lead to a long-lasting change in network input. In these simulations, the average network firing rate was reliably brought back to the target firing rate of 2 Hz by both forms of homeostasis (Figs. 4A-C, black traces). As above, in networks with non-diffusive homeostasis this was achieved by returning the rate of each neuron to the target firing rate regardless of their external input (Fig. 4A, colored traces). In contrast, for networks with diffusive homeostasis, we found that the separability of firing rates of individual groups are maintained according to their input, while the firing rates of all groups were simultaneously reduced so that the average network firing rate again reached the target (Fig. 4B, colored traces). Introducing variability in homeostatic targets for the non-diffusive case, as described previously, did not maintain separability of individual groups as in the diffusive case. Instead, the different groups returned to their mean firing rates that existed before inputs were elevated (Fig. 4C).

The distribution of final firing thresholds explains these differences (Figs. 4D–F). For non-diffusive homeostasis, neurons in the group receiving 10 Hz input had the highest thresholds since they needed to reduce their firing rate the most, followed by the 5 Hz and 2 Hz groups respectively. This led to the final threshold of each neuron reflecting its input. Note that the distribution of firing thresholds is broader in this setup than in (Figs. 3A–B), as a broader range of inputs is given to the network. For a diffusive signal, a neuron’s firing threshold is modulated by the activity of nearby neurons. Since group membership of a neuron is independent of its position, this effect again introduced a correlation between each neuron’s threshold and the mean threshold of the entire network, resulting in a distribution of final thresholds which are less segregated according to their input compared to a network with non-diffusive homeostasis. Thus, firing thresholds in neurons undergoing diffusive homeostasis were more weakly related to their external input. This in turn preserves local firing rate differences in input groups while maintaining constant average network activity. Introducing variable targets for non-diffusive homeostasis caused the thresholds to depend more strongly on their external input, similar to the non-diffusive case.

We could broadly reproduce the distinctions between diffusive and non-diffusive homeostasis in the dynamic mean-field approach by varying *α*. For *α* = 0, modeling non-diffusive homeostasis, we obtained identical firing rates in input groups, as in the recurrent network (Figs. 4G). Note that changing the input of groups of neurons in the recurrent network also affects the activity of neurons with fixed input (Figs. 4B, red traces) due to recurrent connections, an effect that is obviously absent in the dynamic mean-field description. Increasing *α* led to local firing rate differences persisting for longer periods of time. However, these differences eventually decay very slowly, only remaining stable for the case where *α* = 1 (Figs. 4H–I). The reason this occurs is, even after the population activity has quickly reached its homeostatic target, the deviations of the input groups still exert a small homeostatic force when *α <* 1. For example, if *α* = 0.95, there will be a relatively fast change in thresholds as the population activity reaches its target, followed by much slower changes, at 1 − α = 0.05 times the speed (Fig. 4I). This does not happen to the same extent in the spiking network simulations with diffusive homeostasis, as diffusion of NO ensure that deviations from the population activity are directly compensated for by neighboring neurons. Differences persist for 3245 ±440 s, compared with 115 ±6 s and 140 ±40 s for non-diffusive homeostasis with uniform and variable homeostatic targets, respectively (data not shown. ±symbol denotes standard error of the mean of 6 independent network realizations in each case, see Methods). Since we have increased the speed of homeostasis in order to reduce simulation time (see Methods), a more realistic time course of 15 minutes for NO modulation would cause input differences to persist in networks with diffusive homeostasis for many hours to days [5]. Taken together, this shows that diffusive homeostasis can retain input heterogeneity due to the influence of neighboring neurons’ activity on an individual neuron’s firing threshold.

### Population heterogeneity during diffusive homeostasis enables linear network responses

In the simulations shown so far, each neuron received a static input throughout since we were interested in the final network states. We now investigate how these networks respond to fast changes in input; specifically how faithfully each neuron represents its change in input. Since networks with diffusive homeostasis simultaneously maintain constant average network activity and firing rate heterogeneity, we expected that this should allow input modulations to be followed more precisely due to a greater representational capability.

After the network reached steady state under an initial distribution of external inputs, we froze homeostasis so as to simulate changes fast changes in activity, since we assume that homeostasis is not active in these time scales. We then regenerated the external inputs to each neuron from the same distribution presented during homeostasis. This can be thought of as a re-configuration of inputs due to external fluctuations. To best represent such changes in a simple population coding paradigm, each neuron should respond linearly to a change in input; non-linear transformations may lead to an information loss and hence affect neural computations, although this may indeed be desirable in some brain regions. We interpreted the range of changes in input over which this response is linear, or non-saturating, as the range over which homeostasis does not interfere with the network response.

Figs. 5A-C show the change in input rate versus change in output rate of each neuron. A highly nonlinear response was observed in networks with non-diffusive homeostasis, with rectification for large decreases and superlinear responses for large increases in input. This effect was quantified by an *R*^2^ value of 0.57 from a linear regression. Conversely, networks with diffusive homeostasis exhibited a linear response across the entire range of input changes, with an *R*^2^ value of 0.85. Population heterogeneity can also be achieved, as discussed before, by introducing target variability during non-diffusive homeostasis. This yielded a similar non-linear response as in the non-diffusive network with homogeneous targets, with an *R*^2^ value of 0.38.

A consequence of the asymmetry in responses to input changes for networks with non-diffusive homeostasis was that the population rate increases upon regenerating inputs, despite the fact that mean input to the network remained unchanged (Fig. 5D). This did not occur for networks with diffusive homeostasis, suggesting that these networks are more adept at maintaining a target level of activity in conditions where external inputs are dynamic and fast-changing. Crucially, the benefits of a diffusive homeostatic signal can be achieved by a relatively broad range of values for the rate of diffusion, *D*, indicating that the effects we describe are robust to precise parameter choices (Fig. 5E).

This difference in responses to input changes could again be reproduced in the dynamic mean-field approach. This allowed us to characterize population responses across different effective diffusive ranges, using the *R*^2^ value from a linear regression as a measure of response linearity. Fig. 5F shows *R*^2^ values across a range of different input distribution widths, *δ*, as *α* is varied to model different diffusion coefficients (see Methods). This revealed a dependency on *δ*: While values of *α* ∼ 1 exhibited the best response for smaller *δ*, hence cases where the inputs are rather narrow, the optimal *α* decreased as *δ* increased, as well as the overall response linearity. This dependence on input width can be explained by considering the manner in which a population of neurons with a distribution of dynamic ranges span a range of inputs. If this range of inputs is small, then all neurons will span it regardless of their dynamic range (determined by their firing threshold), hence the high values of *R*^2^ for *δ* = 0.1. For an intermediate range of inputs, neurons whose dynamics range is best adapted to the average input are most responsive. This is achieved by increasing *α*. If the range of inputs is very large (*δ* = 1.0), *R*^2^ values are low since the dynamic ranges of the population cannot span the inputs. This effect is stronger at high *α*, as firing thresholds are more correlated, and the dynamics range of most neurons cannot capture the full input variance.

Since connection probability falls off with spatial distance in cortical networks [18], we additionally simulated recurrent networks featuring such connectivity profiles. These networks exhibited qualitatively similar behaviour under diffusive and non-diffusive homeostasis compared to networks without any spatial dependence in connectivity (Supporting Information, Fig. S1).

Overall, these results suggest that networks undergoing diffusive homeostasis are better suited to linearly represent a range of inputs. We investigated this by presenting the networks with time-varying inputs after freezing homeostasis. Groups of excitatory neurons received additional inputs which were randomly and independently generated after fixed time intervals (see Methods). Fig. 6A shows the representation of such a time-varying input pattern (dotted black line) for each network (colored lines).

Networks which have undergone diffusive homeostasis were capable of tracking this input significantly better than their non-diffusive counterparts, as characterized by the RMS error between the network response and input pattern (0.12 for diffusive homeostasis; 0.23 and 0.19 for non-diffusive homeostasis with uniform and variable targets, respectively; Fig. 6B).

We can explore these differences further by constructing a simplified task in which a population of orientation-selective neurons respond to the orientation of a stimulus (see Fig. 6C-F, Methods). This is not intended to represent circuits which perform this task in the brain, but to serve purely as a demonstration of the relative merits of linear and non-linear network responses.

Neurons in the network are randomly assigned a preferred stimulus orientation, independent of their spatial position. A stimulus of a certain orientation can then be presented to the network by varying the external input rates of each neuron, with neurons whose preferred orientation is closest to the stimulus orientation receiving the highest input rate. The stimulus orientation can be decoded from the network by taking the vector average of the stimulus response across all neurons. The orientation of this vector average, or population vector, is the decoded stimulus orientation. Networks with linear responses perform better than those with non-linear responses in decoding stimulus orientation, as measured by the standard deviation of errors in the orientation of the population vector compared to the stimulus orientation (41*◦*, 63*◦* and 72*◦* for diffusive homeostasis, non-diffusive, and non-diffusive with variable targets respectively, Fig. 6G).

### Properties of diffusive homeostasis are conserved in networks with Hebbian plasticity

In the networks described so far, we have used static and uniform synaptic weights for recurrent connections. We next considered whether the observed properties of diffusive homeostasis are altered by the presence of plastic synaptic weights, in particular when Hebbian spike-timing-dependent plasticity (STDP) is introduced (see Methods). Using a standard model of STDP with additive depression and potentiation for all recurrent excitatory synapses, we simulated networks with both STDP and homeostasis active until synaptic weight and firing rate distributions reached a steady state [19]. As before, firing rate distributions were broader in networks with diffusive homeostasis (Fig. 7A). Broad distributions could also be achieved by introducing variability in homeostatic targets. Spiking activity remained asynchronous after STDP, as shown by the distribution of inter-spike intervals (Fig. 7A, inset) [20], and the additive STDP rule led to a bimodal distribution of synaptic weights (Fig. 7B), as previously reported [19]. STDP amplified the differences in response linearity that was observed between homeostatic cases. Inputs to each neuron were regenerated from the same distribution presented during plasticity, and the corresponding change in output rate was compared to the change in input rate, as in Figs. 5A-C. While the response linearity, given by the mean *R*^2^ value, for networks with diffusive homeostasis was 0.16, networks with non-diffusive homeostasis exhibited much lower mean values of 0.01 and 0.02, for uniform and variable homeostatic targets respectively (Fig. 7C). Networks without any homeostasis had a mean value of 0.1. The lower *R*^2^ in all cases compared to networks with static synapses is due to the smaller impact that changes in external input have on these networks, as STDP increases the ratio of recurrent input to external input.

We observed qualitatively similar retention of broad firing rate distributions and response linearity with diffusive homeostasis when a weight-dependent update rule was used (not shown), which has been argued to lead to more realistic weight distributions [21].

## Discussion

Neural homeostasis is commonly thought of as a local process, where neurons individually sense their activity levels and respond with a compensatory change if activity changes. Here we investigated a complementary mechanism, where homeostasis is mediated by a diffusive molecule such as NO that acts as a non-local signal. Using a generic recurrent network model, we show that this form of homeostasis can have unexpected consequences. First, we found that it enables and maintains substantial population heterogeneity in firing rates, similar to that observed experimentally in intact circuits [11], and that input heterogeneities can be preserved in the population activity. Second, the specific form of neural heterogeneity brought about by diffusive homeostasis is particularly suited to support linear network responses over a broad range of inputs. It is important to note that this behavior differs from that of networks where heterogeneity is simply introduced by randomly assigning a different target to each neuron. These results predict that disrupting neural diffusive NO signaling can affect perceptual and cognitive abilities through changes of neural population responses. While other non-diffusive homeostatic mechanisms would continue to stabilize neural activity, the lack of a signal related to the average population activity may disrupt the flexible maintenance of firing rate heterogeneity, and as a result the ability to represent network inputs.

Mean-field analysis revealed that these differences are essentially due to the diffusive messenger providing each neuron with a combination of the average network activity and its own activity as the homeostatic signal. Diffusion of the signal from highly active neurons causes a reduction in the activity of their neighbors, such that firing rates of highly active neurons do not have to be completely reduced in order for the population to achieve a target rate. As a consequence, diffusive homeostasis furnishes a network with an efficient way of flexibly maintaining heterogeneity of firing rates. These effects can also be understood by considering the neural transfer functions, as illustrated in Figs. 8A-B, which provides an intuitive explanation for the differences in firing rate distributions observed under diffusive homeostasis [17, 22]. For non-diffusive homeostasis the transfer function of each neuron is brought to center around its input, leading to a narrowing of the firing rate distribution. Diffusive homeostasis decorrelates the input and threshold of individual neurons, resulting in a population of neurons residing along the entire transfer function. This preserves the non-linear shape of the transfer function, causing broad and heavy-tailed firing rate distributions.

Narrow firing rate distributions are an obvious consequence of local homeostatic processes, as for instance shown recently with homeostasis implemented as local synaptic metaplasticity [14]. This is in apparent conflict with the growing body of experiments documenting broad and heavy-tailed distributions of firing rates in cortex [11]. One could argue that a straightforward explanation is a process, for example genetic or developmental, which randomly assigns neurons heterogeneous homeostatic targets. While we show here that this can result in broader firing rate distributions, we also found that this generally leads to networks with a mismatch between the neural dynamic ranges and input statistics, which in turn limits the responsiveness of the network.

A striking feature of diffusive homeostasis is the lack of requirement for any such distribution of homeostatic targets, as the diffusive signal can be effectively exploited through providing a context for heterogeneity-neurons which maintain a significantly higher firing rate than the rest of the network also synthesize a higher level of the diffusive signal, thus ensuring that their deviation from the average firing rate is counterbalanced by lowering neighboring neurons’ firing rates. This mechanism essentially allows neurons to differ in activity from the population as long as the population as a whole provides some compensation for these deviations. Moreover, this mechanism is compatible with the recent finding that a minority of cells were found to consistently be the most highly active and informative across brain states [23]. While non-diffusive homeostasis would have a disruptive effect on such a ‘preserved minority’ of neurons by reducing their activity towards those of the less active majority, diffusive homeostasis provides a substrate for maintaining their differentiated activity.

A significant distinction between the effects of diffusive and non-diffusive homeostasis appears when network responses to rapidly changing input are considered (Fig. 5). We show that networks with diffusive homeostasis represent input changes more faithfully than those with non-diffusive homeostasis. Saturation of neurons’ responses to large changes are observed in networks without diffusion -this effect is further illustrated in Fig. 8C-D.

Across a spatially homogeneous network, diffusing signals act to effectively shift the transfer function of each neuron towards the average network input, ensuring that neurons are responsive across the entire range of inputs presented to a network. This is in contrast to networks with non-diffusive homeostasis, in which individual neurons are only responsive in a range around their current input. Moreover, the asymmetric response of networks with non-diffusive homeostasis causes the average network activity to increase after fast input changes, while it is constant for a network with diffusive homeostasis (Fig. 5D). The latter case is consistent with observations that mean population firing rates are preserved across novel and familiar environments and across different episodes of slow-wave sleep [24, 25]

Networks with diffusive homeostasis have an improved ability to accurately track time varying inputs (Fig. 6A-B) as a direct consequence of their linear responses. Beneficial effects of neural heterogeneity for population coding have been suggested before [13, 26], but here we find that the broad linear response regime maintained by diffusive homeostasis further improves network performance. This improvement in network performance is also observed in a simplified stimulus orientation decoding task (Fig. 6C-G). Networks with diffusive homeostasis perform better than those with non-diffusive homeostasis when a population vector is constructed from the neural responses in order to decode stimulus orientation (Figure 6G-H). Although there exist alternative methods for decoding stimuli, the population vector has been shown to exhibit performance close to the optimal maximum likelihood procedure for broad tuning, as was used in our example [27].

These distinctions between diffusive and non-diffusive homeostasis are conserved in networks with STDP (Fig. 7). This demonstrates that the limitations of non-diffusive homeostasis in maintaining neural heterogeneity and responsiveness extend beyond the case of static inputs, towards more realistic situations in which neurons receive ongoing and diverse perturbations. Indeed, networks with non-diffusive homeostasis lost almost all sensitivity to external inputs after STDP, while networks with diffusive homeostasis retained this sensitivity (Fig. 7C).

The consequences of diffusive and non-diffusive homeostasis coexisting were also explored, by implementing these mechanisms simultaneously in a single network (not shown). Stable homeostatic activity could be robustly maintained, with the resulting network behaviour depending on the relative timescales of the non-diffusive and diffusive mechanisms. If non-diffusive homeostasis acted faster than diffusive homeostasis, the network exhibited a narrow rate distribution and a low responsiveness to input changes. Conversely, if diffusive homeostasis acted faster than non-diffusive homeostasis, the network exhibited broad firing rate distributions and linear responses to input changes. This is a plausible scenario, as NO modulation of ion channels through occurs in a timescale of 15 minutes [5], while other homeostatic processes which require transcriptional changes occur in a timescale of hours to days [28]. These results reflect what is observed as *α* is varied in the dynamic mean-field analysis, as local and global homeostatic mechanisms are simultaneously active for values in the range 0 *< α <* 1.

It is important to note that modelling HIP as a force acting on the threshold of an integrate-and-fire neuron in order to achieve a target firing rate is a significant simplification. More physiologically realistic descriptions of homeostatic processes reveal the complex relationship between ion channel concentrations and the regulation of a wide range of neural activity [28]. Moreover, a number of previous studies have explored the effects of volume transmission on network dynamics, including its potential in implementing a winner-take-all function [29], the ability of a diffusive signal to reflect the average activity of a group of neurons [30], and the role of another diffusive neurotransmitter, TNF*α*, in epileptogenesis [31]. Here, we add a functional interpretation by exploring its effects on neural heterogeneity and responsiveness within a network.

While NO is involved in a wide variety of neural processes throughout development and learning [32–34], these were ignored throughout for the sake of simplicity and tractability. Nonetheless, the impaired performance of nNOS knock-out mice in cognitive tasks [35] and the prevalence of epilepsy following nNOS inhibition [36] could be linked to diminished homeostatic control of neural excitability. Finally, the outcome of this study is not necessarily confined to NO, and could equally apply to other diffusive neurotransmitters observed in the brain such as hydrogen sulfide and carbon monoxide [37]. Indeed, we conclude that it demonstrates the potential role of diffusive neurotransmitters as an economical and reliable signal of activity across a population of neurons.

## Methods

### Network model

We simulated a spiking network of leaky integrate-and-fire (LIF) model neurons with conductance-based synapses and injected Ornstein-Uhlenbeck noise, as described by

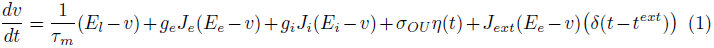

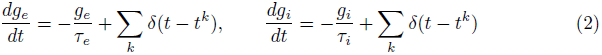

where *v* is the membrane potential, *τ*_*m*_ the membrane time constant, *E*_*l*_ the leak conductance reversal potential, and *σOU* the variance of the injected noise. *η*(*t*) is an Ornstein-Uhlenbeck process with zero mean, unit variance, and correlation time *τ*_*OU*_ = 1 ms [38]. *g*_*e*_ and *g*_*i*_ are the excitatory and inhibitory synaptic currents respectively, given by Equation (2), where *t*_*k*_ denotes the time of all *k* incoming spikes. The reversal potential of the synapses are denoted by *E*_*e*_ and *E*_*i*_, the synaptic conductances by *J*_*e*_ and *J*_*i*_, and the synaptic time constants by *τe* and *τi*. The external input conductance is given by *J*_*ext*_, and *t*^*ext*^ denotes the arrival time of external input, modeled as an independent homogeneous Poisson process for each neuron *i* with rate *µ*_*i*_. A spike is emitted whenever the membrane potential *v* exceeds the firing threshold *θ*, and the membrane potential is then reset to the resting potential value, *v*_*r*_, after a refractory period, *τ*_ref_.

The network was made up of *N* neurons; 0.8*N* excitatory and 0.2*N* inhibitory, with excitatory and inhibitory synaptic conductances scaled so that the network was in a balanced state [16]. Recurrent connections were random and sparse, with connection probability 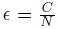 independent of neuron type, where *C* defines the mean number of synapses per neuron. The balanced state was achieved in the network through scaling the inhibitory synaptic conductances by a factor of *g*, such that *J*_*i*_ = _*g*_*J*_*e*_.

### NO synthesis and diffusion

We assumed that neuronal NO synthase (nNOS) is activated by Ca^2+^ influx following a spike (3) and describe the relationship through the Hill equation (4), which results in a sigmoidal concentration dependence, where *n* and *K* are parameters of the Hill equation [39] and *τ*_Ca^2+^_ and *τ*_n_NOS are the timescales of Ca^2+^ decay and nNOS activation respectively.

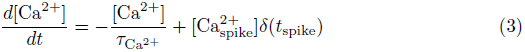

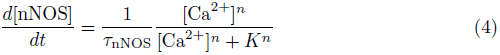

Throughout this paper we considered the case where all neurons, inhibitory and excitatory, express nNOS. The 2D diffusion equation (5) was solved numerically using a spatial resolution *d*_*s*_, with periodic boundary conditions defined by the torus, diffusion coefficient *D* and a decay term *λ* [40]. Neurons were represented by a point source according to their activated nNOS concentration. Periodic boundary conditions were used, as we assume we are simulating a subsection of a cortical network embedded in a larger cortical network with similar network activity.

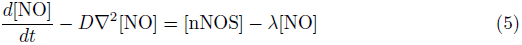

The homeostatic effect of NO was represented in neuron *i* by an increase in *θ*_*i*_, the firing threshold, according to the relative difference in intracellular NO concentration [NO] and a target concentration [NO]_0_;

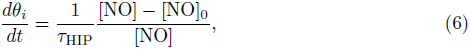

where *τ*_HIP_ is the timescale of homeostasis.

### Non-diffusive homeostasis

For simplicity, the implementation of non-diffusive homeostasis is almost identical to that of diffusive homeostasis, in that the putative non-diffusive neuromodulator [NO_non-diffusive_] is synthesized through equations (3) and (4), and modulates firing thresholds through equation (6). The only difference is that the diffusion term in equation (5) is removed, so that [NO_non-diffusive_] is entirely determined by the rate of intracellular synthesis and decay;

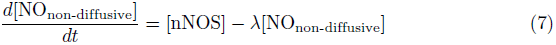

### Dynamic mean-field analysis

For a detailed derivation of equations used in our dynamic mean-field analysis, see [16] and [17]. Briefly, under the assumptions that the network is in an asynchronous regime and that a single EPSP is sufficiently small compared to the voltage required to elicit a spike from resting membrane potential, we can extract the mean firing rate of an LIF neuron in a recurrent network by solving a pair of equations under the condition of self-consistency. The synaptic current for a neuron *i* in a time interval *τ* can be described by its mean *µ*_*i*_ and standard deviation *σ*_*i*_ as follows:

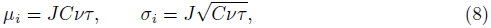

where *J* is the synaptic efficacy, *C* the number of synapses per neuron and *ν* the average population firing rate. The expected mean firing rate *ϕ*_*i*_(*µ*_*i*_, *σ*_*i*_) of an LIF neuron with this synaptic current is given by

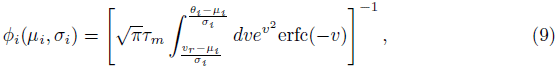

where erfc is the complementary error function. Since the firing rate described by (9) is determined by the synaptic current parameters *µ*_*i*_ and *σi*, which are in turn determined by the population firing rate *ν*, self-consistency requires that the rate which determines the synaptic current parameters must also be equal to the rate which is produced by these parameters, that is:

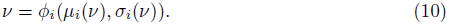

We simulated a non-interacting population of neurons described by the mean-field theory, in which all neurons are identical except for their threshold *θ*_*i*_. Although there is no recurrent excitation within the population, the synaptic current statistics are comparable to that which a neuron within a recurrent network would receive. This enabled us to consider the firing rate distributions arising from presenting single neurons with distributions of synaptic currents, similar to the approach by [17].

We assumed that a neuron embedded in a homogeneous network receiving a diffusive homeostatic signal is analogous to a neuron using a combination of its own firing rate and the average population firing rate as a signal.

The network can then be reduced to a population in which the firing threshold *θ*_*i*_ of each neuron *i* is modulated according to

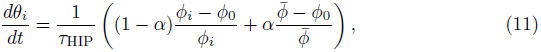

where *ϕ*_0_ is the target firing rate and *ϕ*_*i*_ and 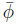 are the firing rate of the neuron *i* and the population respectively. *α* was varied between 0 and 1 and can be thought of as the proportion of NO which a neuron receives due to diffusion from other neurons, with *α* = 0 indicating that each neuron senses only its own activity and *α* = 1 indicating that all neurons share an identical population-wide signal.

In order to implement homeostasis in this setup, we iterated through (8)-(9) until (10) is satisfied to a precision of 10^−4^ Hz, where (9) returns *ϕ*_*i*_ for each neuron in the *ϕ*_*i*_ population, and 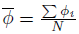 is used as *ν* in (8). At each timestep the thresholds *θ*_*i*_ of each neuron were modulated according to (11), and rates *ϕ*_*i*_ were subsequently recalculated from (9).

### Procedure for Fig. 1

External input rates *µ*_*i*_ for each neuron *i* were randomly drawn from a Gaussian distribution such that 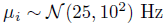 Since the mean NO concentration takes time to reach a steady state in the recurrent network simulations, we ran the network for 100 s without homeostasis and with all neurons receiving 5 Hz input, defining the target NO concentration [NO]_0_ to be the mean NO concentration across all neurons at 100 s.

For the dynamic mean field analysis, we chose parameters which roughly match the rate statistics of the recurrent network simulations. Inputs to each neuron were drawn from a Gaussian distribution such that 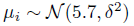, 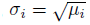. *δ* = 0.4 is the width of the distribution of mean inputs to the population. Note that the parameter *δ* referred to here differs from the *δ* in (1)-(3).

### Adding target variability

In order to match the distribution of effective targets observed during diffusive homeostasis for networks with non-diffusive homeostasis, we assigned each neuron in the non-diffusive network a different homeostatic target, [NO_0_]_*i*_. A network without any homeostasis is presented with input statistics 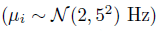, tuned such that the firing rate distribution match that of the network with diffusive homeostasis. [NO_0_]_*i*_ for each neuron *i* can then be drawn from the distribution of steady-state intracellular concentrations of NO for this network. This results in a broad and heavy-tailed distribution of homeostatic targets, as opposed to the single homeostatic target which is used in networks with diffusive homeostasis and the unmodified non-diffusive homeostasis.

A similar approach was adopted in the dynamic mean-field analysis, with each neuron assigned a target firing rate *ϕ*_0,*i*_ from the steady-state firing rate distribution of a network with *α* = 0.8.

### Procedure for Fig. 4

External input for each neuron *i* was *µ*_*i*_ = 2 Hz (*N* = 2500). NO_0_ was set as described previously, although with an input rate of 2 Hz. 2 groups of 250 excitatory neurons each were randomly chosen, independent of neuron position, and stimulated with *µ*_green_ = 5 Hz and *µ*_blue_ = 10 Hz, keeping the inputs to the remaining neurons unchanged. Firing rates plotted in Figs. 4A-C were smoothed with a uniform time window of 20 s. Persistence of input differences were calculated by measuring the length of time it took for the signal-to-noise ratio between the two groups receiving elevated inputs to fall below 0. The signal-to-noise ratio is defined as (*µ*_1_ − *µ*_2_)/(*σ*_1_ + *σ*_2_), where *µ*_*i*_ and *σ*_*i*_ correspond to the mean and standard deviation of the firing rates of group *i*.

### Procedure for Fig. 5

Figs. 5A-D were generated using the same simulation setup as described previously. After the network has reached the homeostatic target firing rate, we froze homeostasis. Input rates to each neuron were then regenerated from the same original input distribution, such that 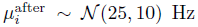. 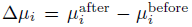 is the difference in input rate each neuron experiences upon this change, and 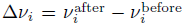 is the corresponding change in output rate for each neuron. The black lines in Figs. 5A-C are from least-squares linear regression, and the *R*^2^ values given were derived from this fit. A similar approach was used in the dynamic mean-field analysis, while varying *δ*, the width of the input distribution.

### Time-varying input

In addition to the external input *µ*_*i*_ previously described, the network was randomly separated into groups of 250 neurons. Each group *j* was stimulated with external Poisson input with a rate given by 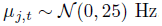. These inputs were regenerated at each timestep *t* of length 1 s. The time-varying input *µ*_*j*,*t*_ was also presented during homeostasis. The dotted black line in Fig. 6A shows the normalized *µ*_*j*,*t*_, while coloured lines show the normalized rate deviation of a randomly chosen group *j* from the mean population firing rate.

### Decoding stimulus orientation

Each excitatory neuron was randomly assigned a preferred orientation. After the network reached a steady state, homeostasis was frozen. For each trial, each neuron *i* with preferred orientation *θ*_*i*_ was stimulated with external Poisson input at a rate given by 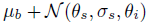, where *µ*_*b*_ = 20 Hz is the base input rate and 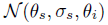 is the amplitude at *θ*_*i*_ of a Gaussian tuning curve centered around the stimulus orientation *θ*_*s*_, with a width given by *σ*_*s*_ = 90^◦^ and a peak amplitude of 2.5 Hz. The angle decoded using the population vector method is the angle of the vector sum of all neural responses.

### Spike-timing-dependent plasticity

A spike-timing dependent plasticity rule, as described in [19], was implemented in each recurrent excitatory-excitatory synapse. Both potentiation and depression are additive in this rule, with no weight dependence. For each pair of pre-and post-synaptic spikes separated by a time ∆*t*, the synaptic weight is updated by a value ∆*w* given by

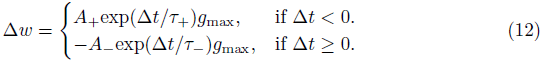

Above, *τ*_+_ and *τ*_*−*_ denote the timecourse over which potentiation and depression occur respectively, while *A*_+_ and *A*_−_ denote the relative strengths of potentiation and depression. *g*max is the maximum synaptic weight. *A*_+_ < *A*_−_ such that irregular firing is maintained within a reasonable range of rates. The external input in this case was given by 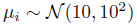, and *J*_ext_ = 40 nS, *C* = 250.

### Model parameters

Unless explicitly defined, the parameters used throughout the paper are given in Table 1. For synthesis, diffusion, and decay of NO we have attempted to match data when available [40, 41], although the dearth of experimental measurements does not permit for great precision [42, 43]. Additionally, parameters were chosen such that the timescale of homeostasis is separated from that of firing rate fluctuations. This is a reasonable assumption, given that activity-dependent NO modulation likely acts within 10 minutes or slower [5], although NO diffusion occurs in the order of 10 seconds. *τ*HIP was chosen to be long enough so as to avoid oscillations but short enough so as to allow feasible large scale simulations. This is a common assumption in computational studies [10]. Larger simulations, up to *N* = 25000, were run with no discernible difference in results.

**Table 1.** Simulation Parameters.

| Neuron |  | NO synthesis and diffusion |  | Network |  | STDP |  |
| --- | --- | --- | --- | --- | --- | --- | --- |
| $E_l$ | -80 mV | $[Ca_{spike}^{2+}]$ | 1 | $N$ | 5000 | $\tau_+$ | 20 ms |
| $v_r$ | -60 mV | $\tau_{Ca^{2+}}$ | 10 ms | $N_e$ | $0.8 N$ | $\tau_-$ | 20 ms |
| $c_m$ | 0.2 nF | $\tau_{nNOS}$ | 100 ms | $J_e$ | 4.0 nS | $A_+$ | 0.025 |
| $\tau_m$ | 20 ms | $n$ | 3 | $J_i$ | 64.0 nS | $A_-$ | 0.0275 |
| $\tau_{ref}$ | 5 ms | $K$ | 3 | $C$ | 100 | $g_{max}$ | 10 nS |
| $\theta_0$ | -50 mV | $D$ | $1000 \mu m^2 s^{-1}$ | $J_{ext}$ | 80.0 nS | | |
| $\tau_e$ | 3 ms | $\lambda$ | $0.1 s^{-1}$ | $\tau_{OU}$ | 1 ms | | |
| $\tau_i$ | 7 ms | Grid size | $1 mm^2$ | $\sigma_{OU}$ | 1 mV | | |
| $E_e$ | 0 mV | $ds$ | $2 \mu m$ | **Dynamic mean-field** | | | |
| $E_i$ | -70 mV | $dt$ (diffusion) | 1 ms | $\theta_0$ | | 10 | |
| $dt$ | 0.1 ms | $\tau_{HIP}$ | 2500 ms | $v_r$ | | 0 | |

All numerical simulations were implemented using the Brian simulator, v1.4.1 [44], and the mean-field analysis was implemented using IPython Notebook [45]. Code and IPython Notebooks which perform the data analysis and plotting will be available on ModelDB and a public github repository following peer review. In the meantime a minimal example of diffusion of a neurotransmitter over a 2D surface is available at http://tinyurl.com/sweeney-diffusion, which may easily be incorporated into existing Brian models.

## Supporting Information

### Procedure for Fig. S1

Spatial dependence in the connection probability between two neurons was introduced as follows:

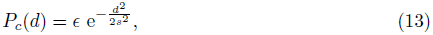

where *d* is the Euclidean distance between the neurons and *s* is a constant defining the connectivity range of the network. 2D positions on the torus are bounded such that *x*, *y* ∈ (0, 1). Given a diffusive range of 0.1, values for *s* in Fig. 1 were therefore set as 0.05, 0.1, and 0.5. The ratio of *s* and the diffusive range was defined as *ρ*, which had values of 0.5, 1.0 and 5.0.

## Acknowledgments

We thank Mark van Rossum, David Sterratt, Jochen Treisch and Alex Cayco Gajic for their careful reading of the manuscript and helpful advice. This work was supported by MRC CDA Fellowship G0900425 (MHH), and the EuroSPIN Erasmus Mundus doctoral programme, UK Engineering and Physical Sciences Research Council, UK Biotechnology and Biological Sciences Research Council and the UK Medical Research Council (grant numbers EP/F500385/1 and BB/F529254/1 for the University of Edinburgh School of Informatics Doctoral Training Centre in Neuroinformatics and Computational Neuroscience) (YS).

---

**Figure 1.**
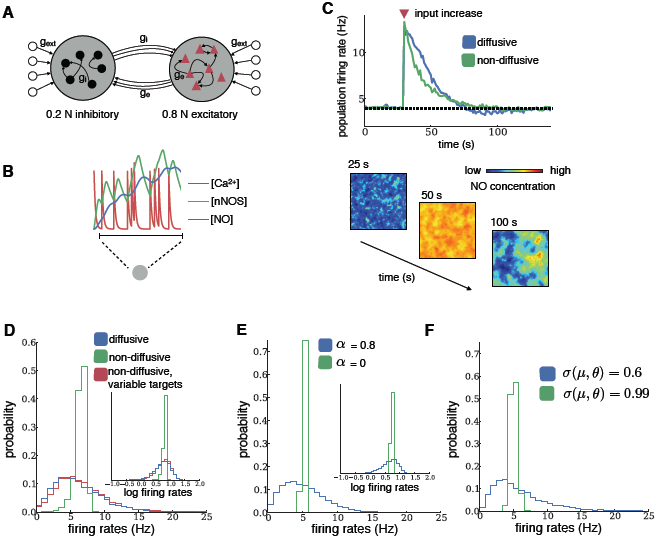
Steady-state behavior of diffusive and non-diffusive homeostasis. **(A)** Schematic of the sparsely connected recurrent network model. Neurons received homogeneous random spiking input (*g*_ext_). **(B)** Intracellular homeostatic signals in a model neuron. Each spike triggers calcium influx, which leads to nNOS activation and NO synthesis. **(C)** Mean population firing rates for networks with diffusive or non-diffusive homeostasis after an increase in external input (red triangle). Spatial distribution of NO concentrations at different times across the network with diffusion are shown below. (**D**-**E**) Distributions of firing rates and log firing rates (insets) after homeostasis from network simulations (D) and mean-field analysis (E), both receiving independent Poisson inputs drawn from a Gaussian distribution. **(F)** Distributions of firing rates in the mean-field analysis for low and high covariance *σ* of threshold and input rate.

**Figure 2.**
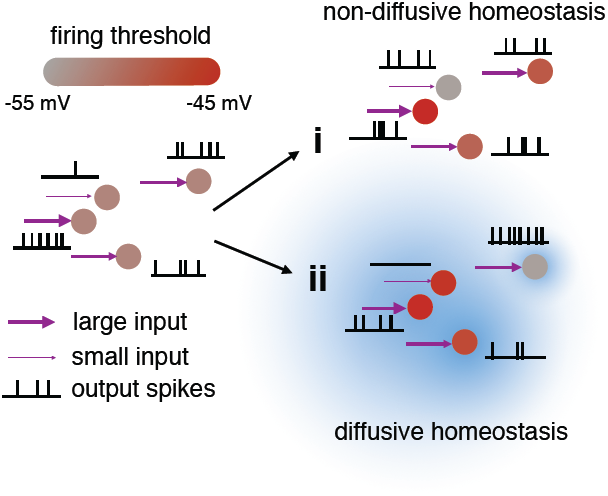
Illustration of the effects of non-diffusive (i) and diffusive (ii) homeostasis. Non-diffusive homeostasis adjusts each neuron’s threshold (red color bar) according to its input to give identical firing rates, while diffusive homeostasis induces correlations (blue cloud) in the thresholds of neighboring neurons, thereby maintaining diverse firing rates.

**Figure 3.**
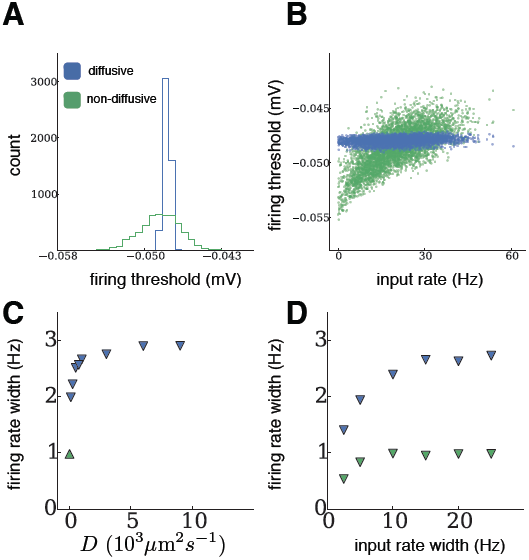
Steady-state firing thresholds of diffusive and non-diffusive homeostasis. **(A)** Distributions of firing thresholds after homeostasis from network simulations, receiving independent Poisson input drawn from a Gaussian distribution. **(B)** Steady-state firing thresholds plotted against external inputs received during homeostasis. **(C)** Width of the steady-state firing rate distribution as the diffusion coefficient *D* is varied (*D* = 1000*µ*m2s*−*1 in panels A-B). **(D)** Width of the steady-state firing rate distribution as the input rate width is varied, for networks with diffusive (*D* = 1000*µ*m^2^s^−1^) and non-diffusive homeostasis (input rate width = 10 Hz in panels A-B).

**Figure 4.**
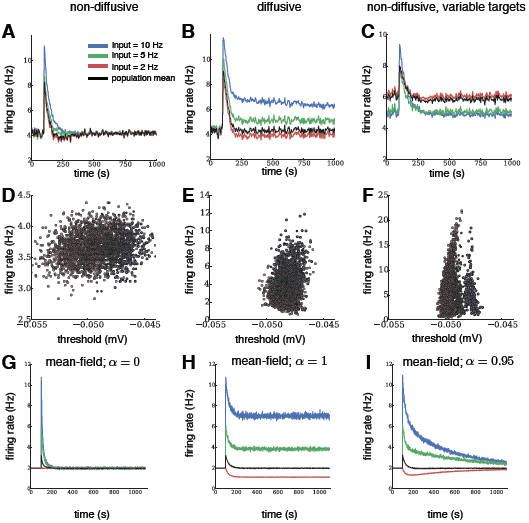
Relative differences between inputs are preserved during diffusive homeostasis. **(A-C)** Evolution of firing rates in a recurrent network for each input group under diffusive and non-diffusive homeostatic control, and for variable homeostatic targets. Black traces show population activity. Independent Poisson input at the target rate was given to 2000 neurons (red), while two groups of 250 neurons each received elevated Poisson input. The relative rate differences of the groups were only preserved for diffusive homeostasis. **(D-F)** Final distributions of firing rates versus firing thresholds. **(G-I)** Evolution of firing rates in a dynamic mean-field population for each input group for purely local (*α* = 0), purely global (*α* = 1), and mixed local and global activity read-out (*α* = 0.95). Black traces show population activity. Independent Poisson input at the target rate is given to 2000 neurons (red), while two groups of 250 neurons each receive elevated Poisson input.

**Figure 5.**
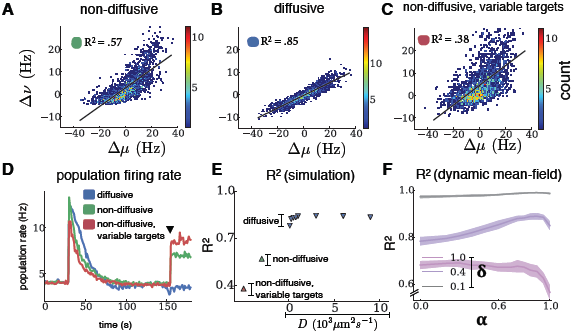
Diffusive homeostasis enables linear network responses. **(A-C)** Firing rate changes ∆*ν* of all neurons following input changes ∆*µ*. Black lines show linear fits, with corresponding *R*^2^ values inset. **(D)** Population activity before and after input change (black triangle). **(E)** Linearity of the population response to a change of inputs as the diffusion coefficient *D* is varied (*D* = 1000*µ*m^2^s^*−*1^ in panel B). **(F)** Linearity of the population response to a change of inputs in the mean-field analysis as *α* is varied, shown for different input distribution widths *δ*. Shaded regions correspond to one standard deviation, averaged over 25 independent network trials for each value of *δ*

**Figure 6.**
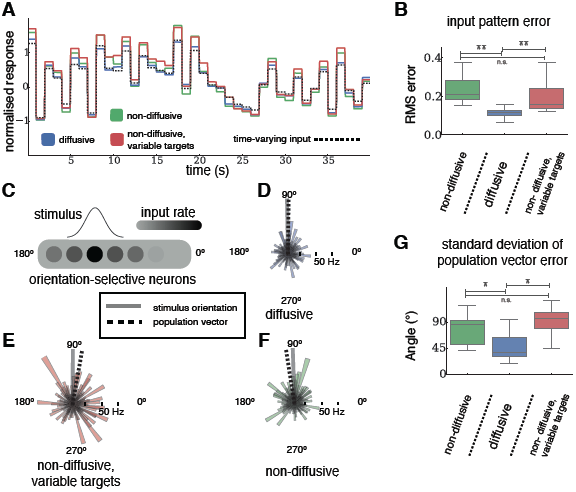
Performance advantages of networks with diffusive over non-diffusive homeostasis. **(A)** Response of each network to a time-varying input pattern (dotted black line). Colored lines show normalized firing rate deviations from the mean population rate of those neurons receiving inputs. **(B)** RMS errors between the normalized input pattern and normalized response of each network (1000 s stimulation and 11 networks in each case). ^∗∗^ *p* < 0.01 from a two-sided Kolmogorov-Smirnov test. **(C)** Diagram showing network inputs during an orientation decoding trial. Each neuron is randomly assigned a preferred orientation, and receives external input at a rate given by the stimulus. **(D-F)** Example response of a population of orientation-selective neurons to a stimulus at an orientation of 90*◦* (grey line), for networks with each type of homeostasis. Individual neural responses and their preferred orientation are given by the radii and direction of the coloured areas respectively. Orientation decoded using the population vector is shown by the dashed line. **(G)** Standard deviation of errors in the orientation of the population vector in response to a stimulus (100 trials for 24 networks in each case). ^∗^ *p* < 0.05 from a two-sided Kolmogorov-Smirnov test. The box encompasses the inter-quartile range and the whiskers extend to 1.5 times the inter-quartile range in all boxplots.

**Figure 7.**
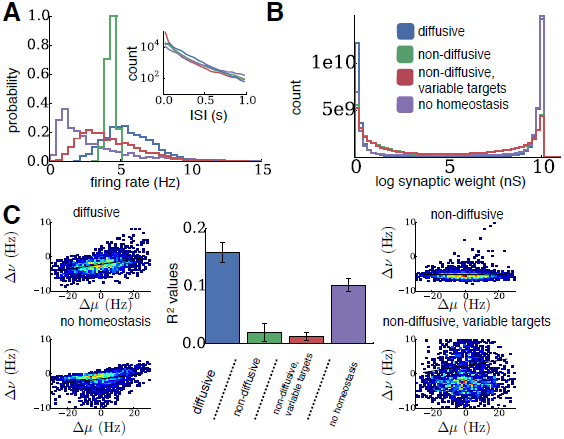
Diffusive homeostasis retains its properties in networks with STDP. **(A)** Distributions of steady-state firing rates after homeostasis and STDP. (Inset) Distribution of inter-spike intervals at the steady-state (log scale). **(B)** Distributions of steady-state synaptic weights after homeostasis and STDP. **(C)** Mean response linearity (measured by *R*^2^ values) for networks with STDP and each case of homeostasis (n=9 networks, error bars represent one standard deviation). Surrounding plots show firing rate changes ∆*ν* of all neurons following input changes ∆*µ* for an example network in each case (black line shows linear fit).

**Figure 8.**
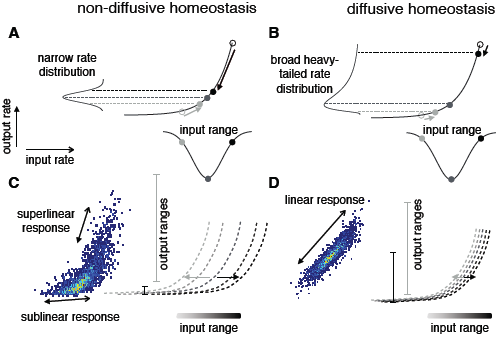
Comparison of the effects of diffusive and non-diffusive homeostasis on neural transfer functions. **(A)** For non-diffusive HIP the transfer function of each neuron is brought to center around its input in order to avoid response saturation or runaway excitation. Arrows show the action of HIP on neurons (circles) with a given input (adapted from [17]). **(B)** Diffusive homeostasis decorrelates the input and threshold of individual neurons, resulting in a population of neurons residing along the entire transfer function. The non-linear shape of the transfer function causes broad and heavy-tailed rate distributions observed with diffusive homeostasis. **(C-D)** Transfer functions following homeostasis of neurons receiving small (light grey) to large (dark grey) input are shown by the dotted lines. Following an instantaneous input change, the output ranges across the entire range of inputs are shown as vertical grey lines, with the shade of grey corresponding to the neuron’s previous input. Networks with non-diffusive homeostasis (C) exhibit response saturation (dark grey line) and superlinear responses (light grey line), while diffusive homeostasis (D) causes transfer functions to shift towards the center of the input distribution, leading to approximately linear responses.

**Figure S1.**
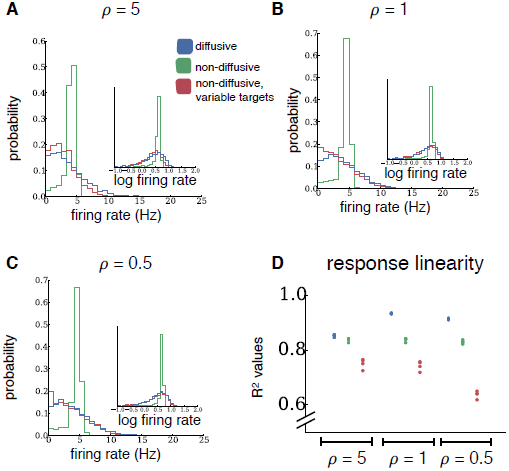
Linearity of network responses in networks with spatially restricted connection probabilities. These results qualitatively agree with those for the random networks used throughout the study. The simulations shown here were identical to those in Figs. 1D and 5 of the main text, but the connection probability between neurons had a Gaussian shape. *ρ*, the ratio of the connectivity range and the diffusive range, is varied across a wide range of values (*ρ*=0.5,1.0,5.0). **(D)** Each point represents the *R*^2^ value of a linear fit as in Figs. 5A-C, for one network.

## References

1. Garthwaite J. Concepts of neural nitric oxide-mediated transmission. Eur J Neurosci. 2008;27(11):2783–2802. Available from: http://dx.doi.org/10.1111/j.1460-9568.2008.06285.x.

2. Pape HC, Mager R. Nitric oxide controls oscillatory activity in thalamocortical neurons. Neuron. 1992;9(3):441–448.

3. Gally J, Montague P, Reeke G, Edelman G. The NO hypothesis: possible effects of a short-lived, rapidly diffusible signal in the development and function of the nervous system. P Natl Acad Sci USA. 1990;87(9):3547–3551. Available from: http://dx.doi.org/10.1073/pnas.87.9.3547.

4. Steinert J, Cornelia K, Baker C, Challiss R, Mistry R, Haustein M, et al. Nitric oxide is a volume transmitter regulating postsynaptic excitability at a glutamatergic synapse. Neuron. 2008;60(4):642–656. Available from: http://dx.doi.org/10.1016/j.neuron.2008.08.025.

5. Steinert J, Robinson S, Tong H, Haustein M, Cornelia K, Forsythe I. Nitric oxide is an activity-dependent regulator of target neuron intrinsic excitability. Neuron. 2011;71(2):291–305. Available from: http://dx.doi.org/10.1016/j.neuron.2011.05.037.

6. Louren¸co CF, Santos RM, Barbosa RM, Cadenas E, Radi R, Laranjinha J. Neurovascular coupling in hippocampus is mediated via diffusion by neuronal-derived nitric oxide. Free Radic Biol Med. 2014;73(0):421–429.

7. LeMasson G, Marder E, Abbott L. Activity-dependent regulation of conductances in model neurons. Science. 1993;259(5103):1915–1917.

8. Lazar A, Pipa G, Triesch J. SORN: a self-organizing recurrent neural network. Front Comput Neurosci. 2009;3:23. Available from: http://dx.doi.org/10.3389/neuro.10.023.2009.

9. Olypher A, Prinz A. Geometry and dynamics of activity-dependent homeostatic regulation in neurons. J Comput Neurosci. 2010;28(3):361–374. Available from: http://dx.doi.org/10.1007/s10827-010-0213-z.

10. Naud´e J, Cessac B, Berry H, Delord B. Effects of cellular homeostatic intrinsic plasticity on dynamical and computational properties of biological recurrent neural networks. J Neurosci. 2013;33(38):15032–15043. Available from: http://dx.doi.org/10.1523/JNEUROSCI.0870-13.2013.

11. Wohrer A, Humphries M, Machens C. Population-wide distributions of neural activity during perceptual decision-making. Prog Neurobiol. 2013;103:156–193. Available from: http://dx.doi.org/10.1093/cercor/bhg100.

12. Marsat G, Maler L. Neural heterogeneity and efficient population codes for communication signals. J Neurophysiol. 2010;104(5):2543–2555.

13. Tripathy SJ, Padmanabhan K, Gerkin RC, Urban NN. Intermediate intrinsic diversity enhances neural population coding. P Natl Acad Sci USA. 2013 May;110(20):8248–53.

14. Zenke F, Hennequin G, Gerstner W. Synaptic plasticity in neural networks needs homeostasis with a fast rate detector. PLo S Comput Biol. 2013 Nov;9(11):e1003330.

15. O’Leary T, Williams AH, Caplan JS, Marder E. Correlations in ion channel expression emerge from homeostatic tuning rules. P Natl Acad Sci USA. 2013;110(28):E2645–E2654.

16. Brunel N. Dynamics of sparsely connected networks of excitatory and inhibitory spiking neurons. J Comput Neurosci. 2000;8(3):183–208. Available from: http://dx.doi.org/10.1016/S0925-2312(00)00179-X.

17. Roxin A, Brunel N, Hansel D, Mongillo G, van Vreeswijk C. On the distribution of firing rates in networks of cortical neurons. J Neurosci. 2011;31(45):16217–16226. Available from: http://dx.doi.org/10.1523/JNEUROSCI.1677-11.2011.

18. Holmgren C, Harkany T, Svennenfors B, Zilberter Y. Pyramidal cell communication within local networks in layer 2/3 of rat neocortex. J Physiol. 2003;551(1):139–153.

19. Song S, Miller KD, Abbott LF. Competitive Hebbian learning through spike-timing-dependent synaptic plasticity. Nat Neurosci. 2000;3(9):919–926.

20. Vogels TP, Abbott LF. Signal propagation and logic gating in networks of integrate-and-fire neurons. The Journal of neuroscience. 2005;25(46):10786–10795.

21. Van Rossum MC, Bi GQ, Turrigiano GG. Stable Hebbian learning from spike timing-dependent plasticity. J Neurosci. 2000;20(23):8812–8821.

22. Ermentrout B. Linearization of FI curves by adaptation. Neural Comput. 1998;10(7):1721–1729.

23. Mizuseki K, Buzsáki G. Preconfigured, Skewed Distribution of Firing Rates in the Hippocampus and Entorhinal Cortex. Cell Reports. 2013;4(5):1010–1021. Available from: http://dx.doi.org/10.1016/j.celrep.2013.07.039.

24. Hirase H, Leinekugel X, Czurkó A, Csicsvari J, Buzsáki G. Firing rates of hippocampal neurons are preserved during subsequent sleep episodes and modified by novel awake experience. P Natl Acad Sci USA. 2001;98(16):9386–9390.

25. Slomowitz E, Styr B, Vertkin I, Milshtein-Parush H, Nelken I, Slutsky M, et al. Interplay between population firing stability and single neuron dynamics in hippocampal networks. eLife. 2015;4.

26. Shamir M, Sompolinsky H. Implications of neuronal diversity on population coding. Neural Comput. 2006;18(8):1951–1986.

27. Seung HS, Sompolinsky H. Simple models for reading neuronal population codes. P Natl Acad Sci USA. 1993;90(22):10749–10753.

28. O’Leary T, Williams AH, Franci A, Marder E. Cell types, network homeostasis, and pathological compensation from a biologically plausible ion channel expression model. Neuron. 2014;82(4):809–821.

29. Kohonen T. Physiological interpretation of the self-organizing map algorithm. Neural Networks. 1993;6(6):895–905.

30. Ott SR, Philippides A, Elphick MR, O’Shea M. Enhanced fidelity of diffusive nitric oxide signalling by the spatial segregation of source and target neurones in the memory centre of an insect brain. European Journal of Neuroscience. 2007;25(1):181–190.

31. Savin C, Triesch J, Meyer-Hermann M. Epileptogenesis due to glia-mediated synaptic scaling. Journal of The Royal Society Interface. 2009;6(37):655–668.

32. Nikonenko I, Nikonenko A, Mendez P, Michurina T, Enikolopov G, Muller D. Nitric oxide mediates local activity-dependent excitatory synapse development. P Natl Acad Sci USA. 2013;110(44):E4142–E4151. Available from: http://dx.doi.org/10.1073/pnas.1311927110.

33. Tamagnini F, Barker G, Warburton C, Burattini C, Aicardi G, Bashir Z. Nitric oxide-dependent LTD but not endocannabinoid-LTP is crucial for visual recognition memory. J Physiol. 2013;591(16):3963–3979. Available from: http://dx.doi.org/10.1113/jphysiol.2013.254862.

34. Son H, Hawkins RD, Martin K, Kiebler M, Huang PL, Fishman MC, et al. Long-term potentiation is reduced in mice that are doubly mutant in endothelial and neuronal nitric oxide synthase. Cell. 1996;87(6):1015–23.

35. Weitzdoerfer R, Hoeger H, Engidawork E, Engelmann M, Singewald N, Lubec G, et al. Neuronal nitric oxide synthase knock-out mice show impaired cognitive performance. Nitric Oxide. 2004;10(3):130–140. Available from: http://dx.doi.org/10.1016/j.niox.2004.03.007.

36. Del-Bel E, Oliveira P, Oliveira J, Mishra P, Jobe P, Garcia-Cairasco N. Anticonvulsant and proconvulsant roles of nitric oxide in experimental epilepsy models. Braz J Med Biol Res. 1997;30(8):971–979.

37. Wang R. Two’s company, three’s a crowd: can H2S be the third endogenous gaseous transmitter? FASEB J. 2002;16(13):1792–1798. Available from: http://dx.doi.org/10.1096/fj.02-0211hyp.

38. Uhlenbeck GE, Ornstein LS. On the theory of the Brownian motion. Phys Rev. 1930;36(5):823.

39. Hill AV. The possible effects of the aggregation of the molecules of haemoglobin on its dissociation curves. J Physiol. 1910;40:4–7.

40. Philippides A, Husbands P, O’Shea M. Four-dimensional neuronal signaling by nitric oxide: a computational analysis. J Neurosci. 2000;20(3):1199–1207.

41. Salerno JC, Ghosh DK. Space, time and nitric oxide–neuronal nitric oxide synthase generates signal pulses. FEBS journal. 2009;276(22):6677–6688.

42. Hall C, Garthwaite J. What is the real physiological NO concentration in vivo? Nitric Oxide. 2009;21(2):92–9103. Available from: http://dx.doi.org/10.1016/j.niox.2009.07.002.

43. Batchelor AM, Bartus K, Reynell C, Constantinou S, Halvey EJ, Held KF, et al. Exquisite sensitivity to subsecond, picomolar nitric oxide transients conferred on cells by guanylyl cyclase-coupled receptors. P Natl Acad Sci USA. 2010 Dec;107(51):22060–5.

44. Goodman D, Brette R. Brian: a simulator for spiking neural networks in python. Front Neuroinform. 2008;2:192–7. Available from: http://dx.doi.org/10.3389/neuro.11.005.2008.

45. Pérez F, Granger BE. IPython: a System for Interactive Scientific Computing. Comput Sci Eng. 2007 May;9(3):21–29. Available from: http://ipython.org.

